# Detecting Operons in Bacterial Genomes via Visual Representation Learning

**DOI:** 10.1101/860221

**Authors:** Rida Assaf, Fangfang Xia, Rick Stevens

## Abstract

Contiguous genes in prokaryotes are often arranged into operons. Detecting operons plays a critical role in inferring gene functionality and regulatory networks. Human experts annotate operons by visually inspecting gene neighborhoods across pileups of related genomes. These visual representations capture the inter-genic distance, strand direction, gene size, functional relatedness, and gene neighborhood conservation, which are the most prominent operon features mentioned in the literature. By studying these features, an expert can then decide whether a genomic region is part of an operon. We propose a deep learning based method named Operon Hunter that uses visual representations of genomic fragments to make operon predictions. Using transfer learning and data augmentation techniques facilitates leveraging the powerful neural networks trained on image datasets by re-training them on a more limited dataset of extensively validated operons. Our method outperforms the previously reported state-of-the-art tools, especially when it comes to predicting full operons and their boundaries accurately. Furthermore, our approach makes it possible to visually identify the features influencing the network’s decisions to be subsequently cross-checked by human experts.

## 1 Introduction

Genes in prokaryotic genomes assemble in clusters, forming transcription units called operons. These genes share a common promoter and terminator^1^, and are usually metabolically or functionally related. Predicting operons helps understand high level organization of genes and regulatory networks^2–8^, annotate gene functions^9^, develop drug candidates^10^, and inhibit antibiotic resistance^11^. While operons are prevalent in bacterial genomes, their detection is challenged by a multitude of factors that contribute to their organization.

Human experts annotate operons by visually inspecting stretches of genes in a comparative genomics browser (see an example in Figure 1). Such visual representations enable synthesis of two sources of information. First, gene-level features such as size, strand direction, function label, and inter-genic distance can be checked for consistency by scanning contiguous regions within the same genome. Second, close relatives of the query genome can be retrieved, aligned, and anchored against the focus gene, which allows human experts to see whether there is evolutionary evidence of regional conservation. This second dimension of criteria is critical to the operon call, but is often difficult to quantify, requiring human judgment on selecting phylogenetic distance and weighing the similarity of genomic neighborhoods.

**Figure 1.**
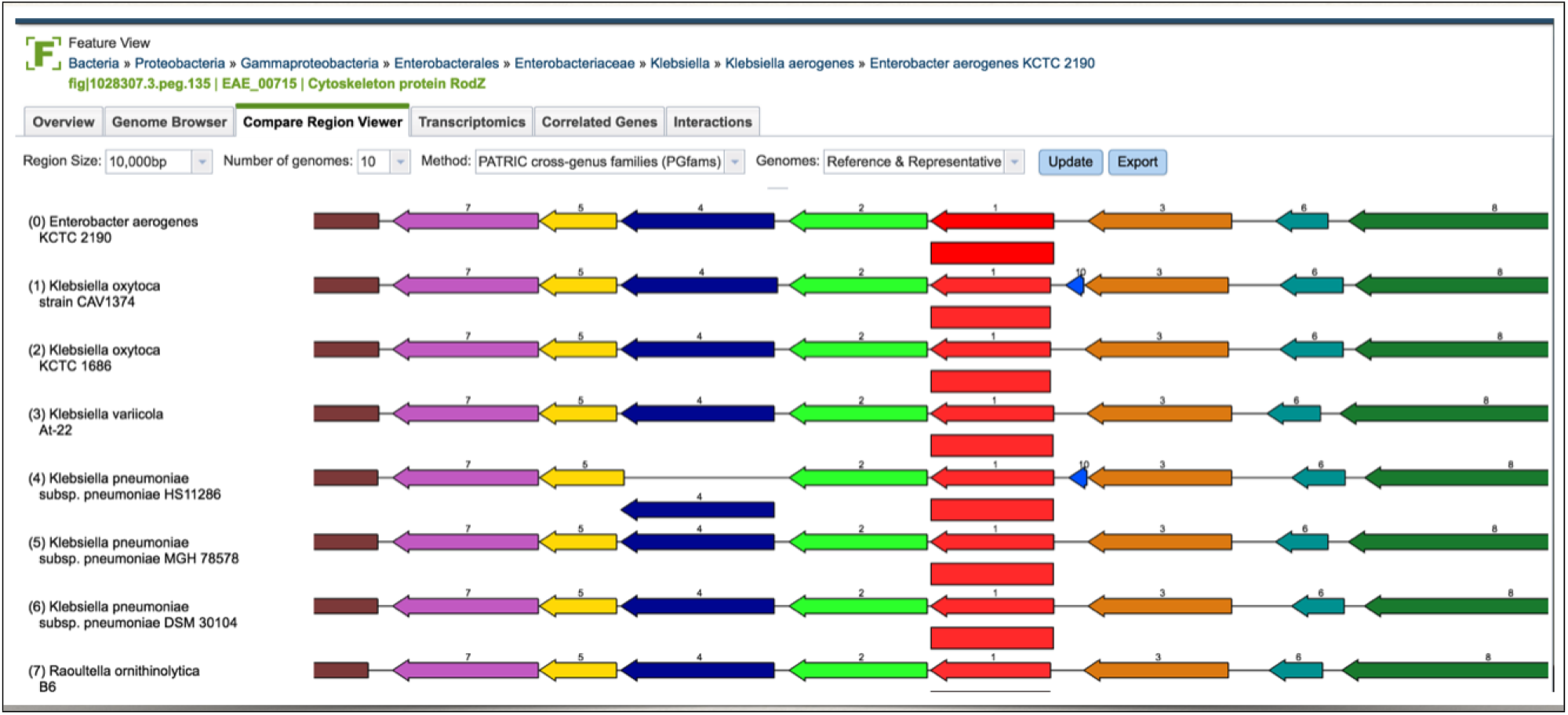
Snapshot of the Compare Region Viewer service provided by PATRIC (https://www.patricbrc.org). The image shows a genomic region of the query genome (first row) aligned against a set of other genomes, anchored at the focus gene (represented as a red arrow). The service starts with finding other genes that are of the same family as the focus gene, and then aligns their flanking regions accordingly.

Several tools have been proposed to detect operons computationally, yet they mostly focus on gene-level features. Even when phylogenetic information is added, it’s preprocessed and presented in a tabular form, preventing flexible learning. We are inspired by how human experts work and the recent advance in deep learning to explore a novel approach based on visual learning. We hypothesize that, when presented with all evidence in a similarly visual representation of genomic images that humans rely on, neural networks can learn to capture the complex operon determinants within and across genomes. To this end, we present a method, named Operon Hunter, that predicts operons from visual representations of genomic fragments. Our method uses a pre-trained network via transfer learning to leverage the power of deep neural networks trained on image datasets. The network is re-trained on a limited dataset of extensively validated and experimentally verified operons. We compare our method with the state-of-the-art operon predictors. Our results show that Operon Hunter outperforms them in identifying full operons as well as delineating operon boundaries. Furthermore, our visual approach generates insights into regions of importance that can be cross-checked by human experts.

### 1.1 Related Work

Different methods focus on different operon features to make their predictions. Some methods use Hidden Markov Models (HMMs) to find shared promoters and terminators^12–14^. Other methods rely on gene conservation information^15^, while others leverage functional relatedness between the genes^2,16^. The most prominent features used in operon prediction are transcription direction and inter-genic distances, as reported in the literature^2,4,17–27^. Gene conservation is another important feature, since adjacent genes that are co-transcribed are likely to be conserved across multiple genomes^6,28^. Different machine learning (ML) methods are used to predict operons, such as neural networks^17,18^, support vector machines^19^, and decision tree-based classifiers^20^. Other tools utilize Bayesian probabilities^21–23^, genetic algorithms^24^, and graph-theoretic techniques^16,22^.

We focus our attention on two machine learning based tools. The first tool^29^ developed by Zaidi and Zhang is reported to have the highest accuracy among operon prediction methods. It is based on an artificial neural network that uses inter-genic distance and protein functional relationships^4^. To infer functional relatedness, this method uses the scores reported in the STRING Database^30^. The STRING database captures functional relatedness through scores generated using information about gene neighborhood, fusion, co-occurrence, co-expression, protein-protein interactions, in addition to information extracted by automatic literature mining. The predictions made by this method were compiled into what is called the Prokaryotic Operon Database (ProOpDB)^5^, and released later as a web service called Operon Mapper^31^. For simplicity, we will refer to this method as ProOpDB, given that it was our resource for the predictions over the model organisms we used for the cross-tool comparison. The second tool is called the Database of Prokaryotic Operons (Door)^3^ and was ranked as the second best operon predictor after ProOpDB by the same study^29^. It was also ranked as the best operon predictor among 14 tools by another independent study^32^. Door’s algorithm uses a combination of a non-linear (decision-tree based) classifier and a linear (logistic function-based classifier) depending on the number of experimentally validated operons available for the genomes used in the training process. Among the features Door uses to perform its predictions for adjacent gene pairs are: the distance between the two genes, the presence of a specific DNA motif in the genomic region separating them, the ratio of the genes’ lengths, the genes’ functional similarity (determined using Gene Ontology (GO)), and the level of conservation of the genes’ neighborhood.

## 2 Results

Most operon prediction tools make their predictions on isolate gene-pairs. Contiguous gene pairs predicted to be as part of an operon are then aggregated to form full operon predictions^2^. We refer to gene pairs that are part of an operon as operonic, and gene pairs where one is a boundary of an operon, or that include a verified operon consisting of a single gene, to be non-operonic. Our validation dataset consists of two well-studied genomes with experimentally validated operons: E. coli and B. subtilis. These genomes are extensively used to verify the majority of the published tools’ results.We compare the performance of the tools first when considering gene-pair predictions, and then considering full-operon predictions. While Operon Hunter demonstrates an advantage in both, the advantage is more pronounced when assessing the ability to predict full operons and their boundaries accurately.

### 2.1 Gene-Pair Prediction

We report the sensitivity (true positive rate), precision, and specificity (true negative rate) for the models’ performance over gene pair predictions in Table 1. These statistical measures were calculated according to the following definitions:

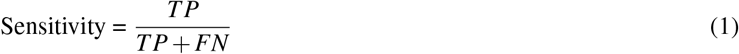

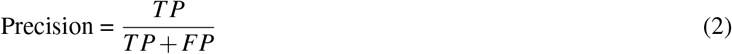

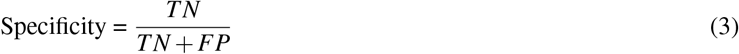

**Table 1.** Sensitivity, Precision, and Specificity. Sensitivity (true positive rate) is the percentage of operonic gene pairs that were detected by the different tools, precision is the percentage of operonic gene pairs predicted by the different tools which are actually true positives, and specificity (true negative rate) is the percentage of Non-operonic gene pairs that were detected by the different tools. For sensitivity and specificity, results are first shown per genome, and then as an aggregate over the entire testing dataset.

|  | Sensitivity<br>E. coli<br>361 pairs | Sensitivity<br>B. subtilis<br>369 pairs | Sensitivity<br>Aggregate<br>730 pairs | Precision<br>Aggregate<br>730 pairs | Specificity<br>E. coli<br>461 pairs | Specificity<br>B. subtilis<br>302 pairs | Specificity<br>Aggregate<br>763 pairs |
| --- | --- | --- | --- | --- | --- | --- | --- |
| Operon Hunter | 88% | 97% | 92% | 92% | 95% | 88% | 92% |
| ProOpDB | 93% | 93% | 93% | 89% | 90% | 88% | 89% |
| Door | 81% | 86% | 84% | 94% | 94% | 97% | 95% |

Where TP is the number of operonic pairs predicted correctly (true positives), FP is the number of non-operonic pairs incorrectly predicted as operonic (false positives), FN is the number of operonic pairs incorrectly predicted as non-operonic (false negatives), and TN is the number of non-operonic pairs predicted correctly (true negatives).

ProOpDB scores the highest sensitivity but the lowest precision. The opposite is true for Door, which achieves the lowest sensitivity but the highest precision. Operon Hunter’s performance is more stable across the two metrics. To capture the predictive power of the model on both classes in a single metric, we report the F1 score, accuracy, and the Mathews Correlation Coefficient (MCC) in Table 2, calculated using the following definitions:

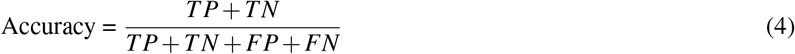

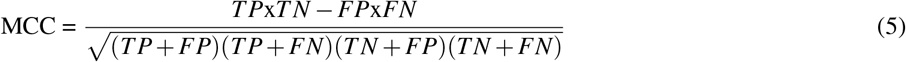

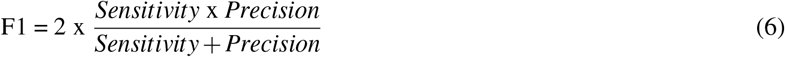

**Figure 2.**
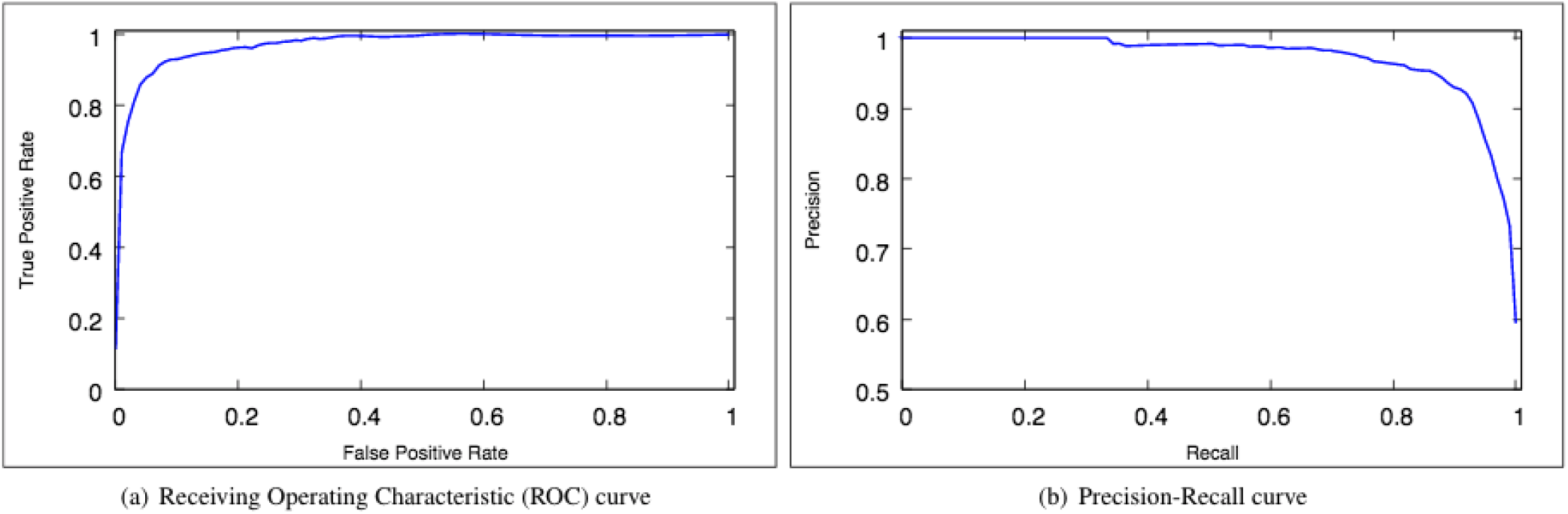
The Receiving Operating Characteristic (ROC) curve (a) and Precision-Recall curve (b) corresponding to Operon Hunter evaluated over the entire testing dataset. The Area Under the Curve (AUC) for both is 0.97.

Operon Hunter scores the highest on all metrics measured, followed by ProOpDB then Door. We also show the Receiver Operating Characteristic (ROC) curve and the Precision-Recall curve with the corresponding Area Under the Curve (AUC) in Figure 2. Evaluated on the entire testing dataset, Operon Hunter scores a ROC AUC and a Precision-Recall AUC of 0.97.

**Table 2.** Accuracy, MCC (Mathews Correlation Coefficient), and F1-score achieved by the different tools over the entire testing dataset.

| Tool | Operon Hunter | ProOpDB | Door |
| --- | --- | --- | --- |
| Accuracy | 92% | 91% | 89% |
| MCC | 0.84 | 0.82 | 0.79 |
| F1-score | 0.921 | 0.908 | 0.886 |

### 2.2 Operon Prediction

Predicting full operons is a more challenging task that requires the accurate identification of the operon endpoints. We reserve the definition of operons as clusters including at least two genes. Matching earlier reports concerning different methods^29^, the accuracies of all 3 tools drop when validated on full operon predictions rather than separate predictions made for every gene pair. We present the percentage of operons predicted fully by each of the tools in Table 3. Full matches are operons reported in the literature that are accurately predicted with the same gene boundaries as the published endpoints. We only consider operons that consist of more than one gene, and were reported in ODB in addition to RegulonDB or DBTBS. This amounted to 254 full operons. Predictions that only partially match a verified operon are not shown. To make full operon predictions, the model starts by generating a prediction for every consecutive gene pair in a genome. In a second pass, pairs predicted as operonic are then merged into full operons. Operon Hunter accurately identifies 85% of the operons fully, which is the most across the 3 tools. Both ProOpDB and Door show a drop in accuracy, to 62% and 61% respectively.

**Table 3.** Comparison of the results between OperonHunter, ProOpDB, and Door when considering full operon predictions. Exact Operon Matches are the percentage of operons predicted where the endpoints exactly match those of the experimentally verified operons. The percentages are reported over 254 full operons that consist of more than one gene and are reported in RegulonDB/DBTBS and ODB.

|  | Operon Hunter | ProOpDB | Door |
| --- | --- | --- | --- |
| Exact Operon Matches | 85% | 62% | 61% |

### 2.3 Cross Validating Visual and Operon Features

An advantage of visual models is that they can generate insights into decision making by highlighting regions of importance. Such interpretable representations reveal inner workings of the model and can be checked by experts to see if they ground their intuition. To investigate the model’s performance, we used the Grad-Cam^33^ method to overlay heat-maps over the input images, highlighting the areas of attention that most affect the network’s decisions. Figure 3 shows some of the network’s confident predictions.

**Figure 3.**
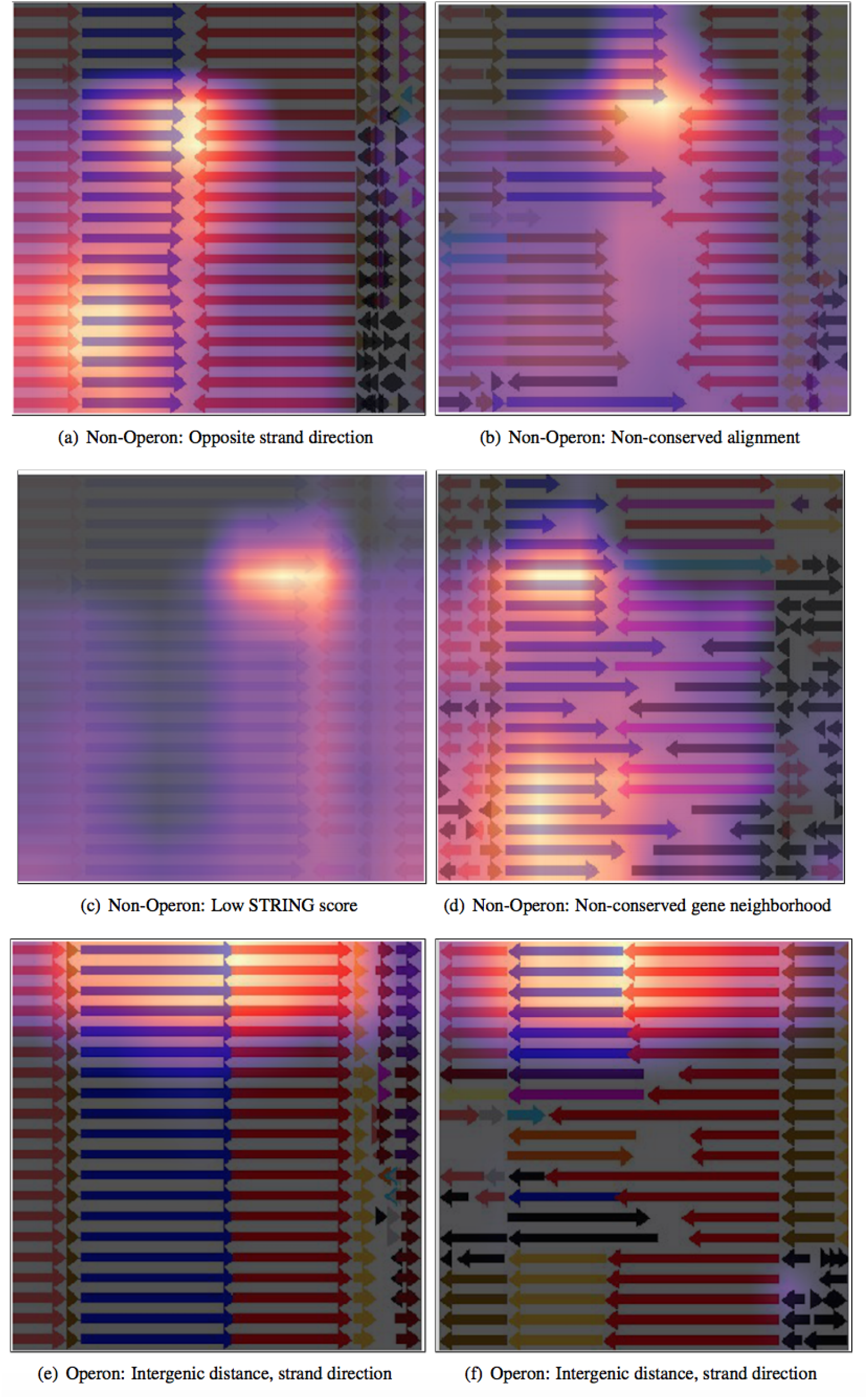
HeatMaps generated using the Grad-Cam method and overlayed over input images, highlighting the network’s areas of attention that had most influence over the network’s decision. Each sub-figure shows the network’s correctly predicted label, with what we believe to be the most prominent feature leading to the network’s decision.

It appears that when predicting a gene pair as non-operonic, the model bases its decision on the entirety of the input image, whereas it focuses on the immediate vicinity of the gene pairs predicted as operonic. Upon closer inspection, we speculate that the following are the most important features influencing the model’s decision:

- Figure 3 (a): The strand directionality of the gene pair.
- Figure 3 (b): The mis-alignment between the genomes.
- Figure 3 (c): The low alpha channel of the image, which is a representation of the low STRING score between the query gene pair.
- Figure 3 (d): The non-conservation of the query gene pair neighborhood.

All these features were previously mentioned as among the most prominent in operon identification. Operonic gene pairs show more overall conformity across the input image, with flanking regions being mostly aligned/conserved, and query gene pairs having little or no inter-genic distance, and usually similar directions, as shown in Figure 3 (e). Even when the neighborhood of the query gene pair is not conserved, as shown in Figure 3 (f), the network seems to base its decision on the similar strand direction and lack of an inter-genic distance between the gene pair. Thus, it seems that when predicting gene pairs as operonic, the network focuses on the immediate vicinity of the pair, disregarding other areas of the image. However, the network seems to pick up on anomalies around the gene pair, taking the entirety of the image into consideration when predicting non-operonic images.

## 3 Methods

### 3.1 Datasets

We used the ‘‘Known Operons” section of the Operon DataBase (ODB)^34^ to construct the training dataset. The Known Operons section contains a list of operons that are experimentally verified, which we used to label the generated images. Out of the entire dataset, the six genomes (Table 4) with the most number of labeled operons were selected. These genomes have significantly more operons than the other genomes listed in this section of the database. For each of the selected genomes, our program produces an image for every consecutive pair of genes that are part of a validated operon. To generate the images, each of these genomes is aligned with the set of reference+representative genomes found on PATRIC^35^. For every aligned gene, an image is generated to capture the surrounding 5 kilo base pairs (Kbp) flanking region. The resulting dataset consisted of 4,306 images of operonic gene pairs. To generate the dataset representing the non-operonic gene pairs, we used the standard approach reported by ProOpDB^5^ as follows: Genes that are at the boundaries of known operons are labeleled along with the respective upstream or downstream gene that is not part of that operon as a non-operonic gene pair. We skipped single-gene operons from the training dataset. A balanced dataset was then curated from the total set of images created.

**Table 4.** Breakdown of the training dataset: The genome names and the corresponding number of gene pairs used. Operon pairs were harvested from the Known Operons section of ODB.

| Genome name | Operon pairs | non-Operon pairs |
| --- | --- | --- |
| Escherichia coli | 1,443 | 1,322 |
| Listeria monocytogenes | 806 | 780 |
| Legionella pneumophila | 611 | 791 |
| Corynebacterium glutamicum | 525 | 396 |
| Photobacterium profundum | 447 | 544 |
| Bacillus subtilis | 474 | 457 |
| Total | 4,306 | 4,290 |

### 3.2 Feature Encoding via Visual Representation

We replicated the Compare Region Viewer service offered by PATRIC (The Pathosystems Resource Integration Center), which is a bacterial Bioinformatics resource center that we are part of (https://www.patricbrc.org), by implementing an offline version forked from the production UI. An offline version is necessary for computational efficiency and for consistency in the face of any future UI changes. Figure 4 shows an example of the generated images.

**Figure 4.**
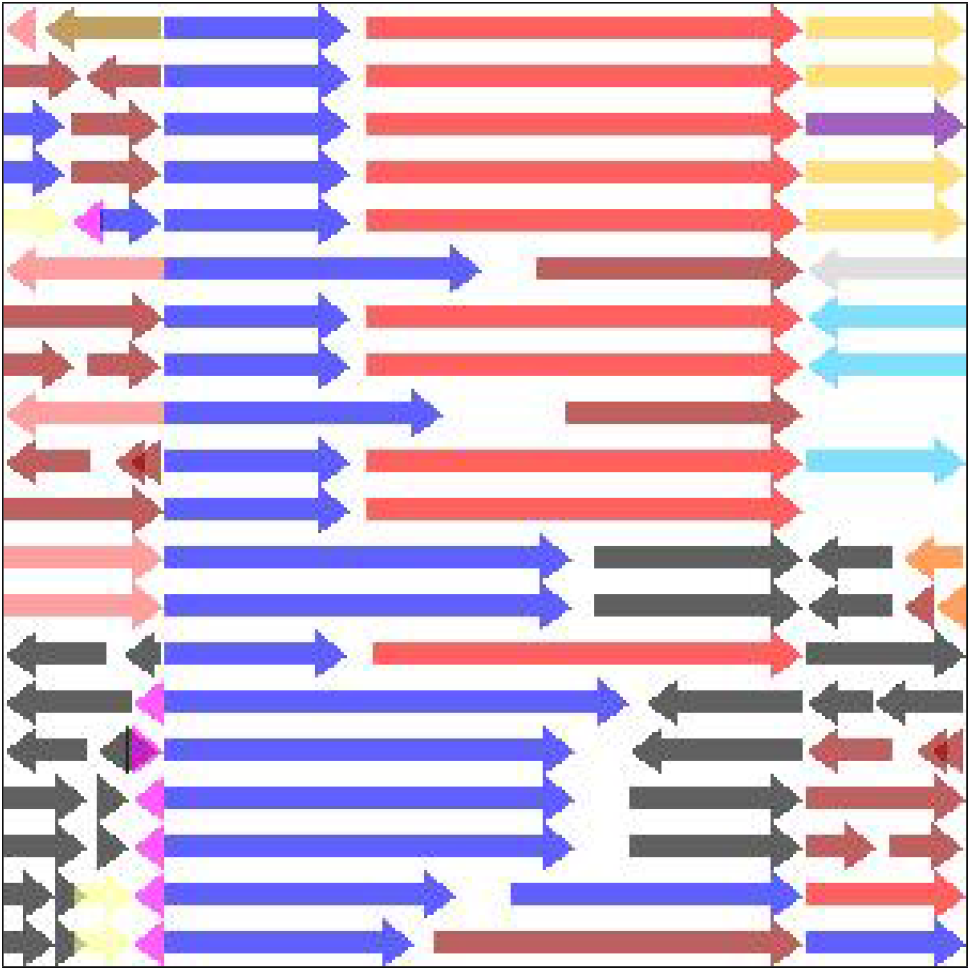
Example of an image generated by our offline version of the Compare Region Viewer service to be fed as input to the neural network. Each arrow represents a single gene. Each row captures the area of interest in a genome. The query genome is the top row. The rest of the rows are genomes selected by evolutionary distance. The query gene pair are colored blue and red. Genes share the same color if they belong to the same family and that family. The query gene pair are centered in the middle, occupying 2/3 of the image’s width. The rest of the flanking region is represented correspondingly to the left/right of the center region. The alpha channel of the image is the STRING score of the query gene pair.

Each row of arrows in the generated image represents a region in a genome, with the query genome being the top row. Each arrow represents a single gene, scaled to reflect its size relative to its region, in addition to the gene’s strand directionality. The distances between the genes are also scaled on each row relative to the gene’s region. Each image consists of three regions. The central region makes up two thirds of the image, and represents the query gene pair. Genes that fall before the query gene pair are represented to the left of the central region, and those that fall after the query gene pair are represented to the right of the central region. Dividing the image implicitly into three regions highlights the area of most interest by zooming in on the query gene pair, while preserving the relevant conservation information of the flanking genomic fragments on the sides. Colors represent gene functionality. The blue and red arrows are reserved to represent the query gene pair and the rest of the genes that belong to the same families. In general, genes share the same color if they belong to the same family, or are colored black after a certain color distribution threshold. The families used in the coloring process are PATRIC’s Global Pattyfams that are generated by mapping signature k-mers to protein functionality, using non-redundant protein databases built per genus before being transferred across genera^36^. Finally, the image’s alpha channel is set to be equal to the STRING score of the query gene pair. If no score exists, the default alpha channel is set to a minimum of 0.1. The generated images capture most of the prominent features mentioned earlier, such as gene conservation, functionality, strand direction, size, and inter-genic distance.

### 3.3 Transfer Learning

The relatively small size of the dataset combined with the depth of the network pose the risk of over-fitting. One way to get around this issue is by using a technique referred to as transfer learning^37^. In transfer learning, a model does not have to be trained from scratch. Instead, a model that has been previously trained on a related task is retrained on the new task. The newly retrained model should then be able to transfer its existing knowledge and apply it to the new task. This approach makes it possible to be able to reuse models that have been trained on large datasets, by adding the necessary adjustments to make them available to work with more limited datasets. This adds a further advantage to representing the data visually. To train and test our model, we used the FastAI^38^ platform. The best performance was observed using the ResNet18 model. All available models were previously trained on Imagenet, which is a database that contains more than a million images belonging to more than a thousands categories^39^. Thus, a model that was previously trained on ImageNet is already good at feature extraction and visual recognition. To make the model compatible with the new task, the top layer of the network is retrained on the operon dataset, while the rest of the network is usually left intact. This is more powerful than starting with a deep network with random weights. Another technique that is commonly used against over-fitting is data augmentation, whereby the training dataset is enriched with new examples by applying certain transforms on the existing images. These transforms include flips (horizontal and vertical), zooming effects, warping, rotation, and lighting changes. Using FastAI’s toolkit, we augmented our training dataset by allowing only one transform, the horizontal flip, to be applied. Given the deliberate feature engineering processes taken to create the input images, we believe that horizontal flips could safely be applied without altering the true nature of the images. This would keep the key information intact and in place (e.g. by keeping the query genome as the top row).

### 3.4 Model Validation

We resort to two extensively studied genomes with experimentally verified operons: E. coli and B. subtilis. These genomes are the standard for verifying operon prediction results in the literature. We limit our validation by using only the experimentally validated operons found in these genomes. We compare the predictions made by Operon Hunter to those made by ProOpDB and Door, the tools with state of the art accuracies as reported by independent studies^3,5,29^. To build the testing dataset (Table 5), we used the operons published in RegulonDB^40^ and DBTBS^41^. RegulonDB is a database containing verified operons found in E. coli, and DBTBS is a database that contains verified operons found in B. subtilis. We cross check the operons found in these databases, with the ones published in the Known Operons section of ODB. We selected the operons that are reported in both: The database corresponding to its organism (RegulonDB for E. coli, DBTBS for B. subtilis), and ODB. The resulting dataset consists of 730 operonic gene pairs. We used the same approach mentioned earlier to construct a non-operonic dataset. Namely, the boundaries of the operons were selected along with their neighboring genes as non-operonic pairs. To construct the non-operonic datasets, it was enough for an operon to be published in any of the mentioned databases to be considered, resulting in a slightly larger non-operon dataset. Using operons that were experimentally verified and published in multiple independent databases adds confidence to the assigned label.

**Table 5.** Breakdown of the testing dataset: The genome names and the corresponding number of gene pairs used. Operon pairs were scoured from RegulonDB (for E. coli) and DBTBS (for B. subtilis) and matched with the Known Operons section of ODB.

| Genome | Operon pairs | non-Operon pairs |
| --- | --- | --- |
| *Escherichia coli* str. K-12 substr. MG1655 | 361 | 461 |
| *Bacillus subtilis* subsp. *subtilis* str. 168 | 369 | 302 |
| Total | 730 | 763 |

Some tools train on the same organisms they report their predictions on. In our experiments, we exclude the testing dataset from the training dataset, by following a leave-one-out approach on the genome level. Thus, none of the genomes used in the training process belong to the same organism as the genome used for testing our method’s performance. For example, when evaluating the model’s predictions on the E. coli dataset, it is trained on all the images representing all the genomes except E. coli, and then tested on the images representing the E. coli genome. This approach leads to an 87-13 train-test split for the E. coli genome, and a 92-8 train-test split for the B. subtilis genome.

## 4 Discussion

We have presented a novel approach to operon prediction by training a deep learning model on images of comparative genomic regions. This approach has achieved better results than the currently available state-of-the-art operon prediction tools. In particular, it has demonstrated a clear advantage in predicting accurate operon boundaries (i.e. predicting operons fully).

The first advantage of our model is in its effective synthesis of gene-level and phylogeny-level evidences. Traditional models come with preprocessed features and may be rigid in picking parameters on neighborhood size, genomic distance, or similarity metrics for comparing regions across genomes. The PATRIC compare region viewer is perfected by human experts, and strikes a balance in diversity and granularity in the way it brings representative genome relatives to an annotator’s attention. Critically, these images give machine learning models a two-dimensional view of all relevant information without limiting them to a pre-determined way of encoding features. High-capacity neural network models are thus allowed a more flexible space to learn and pick up interactive features. For example, the images offer a natural way to compare genes (horizontally) and clusters across genomes (vertically) with 2D convolution. The fact that genomes are sorted by evolutionary distance allows the neural network to exploit locality and place more emphasis on close genomes via incremental pooling.

The second advantage of our model comes from leveraging neural networks pre-trained with image recognition capabilities. Many deep learning model architectures were first developed in vision tasks and perfected over years of iterations. Google researchers have used spectrogram (instead of text) in direct speech translation^42^ and DNA sequence pile-up graphs (instead of alignment data) in genetic variant calling^43^. In both cases, the image-based models outperformed their respective previous state-of-the-art method based on traditional domain features. Further, the low-level visual feature patterns learned in pre-trained image models have been demonstrated to transfer to distant learning tasks on non-image data in several preliminary studies ranging from environmental sound classification to cancer gene expression typing^44^.

We experienced technical difficulties when trying to retrieve the operon predictions made by both tools we are comparing our performance against. Since the tools generate predictions for gene pairs, the shortest possible operon should thus consist of at least two genes. However, this conflicts with the reported predictions made by these tools, as they include operons consisting of only a single gene. We considered single-gene predictions to be part of a non-operonic gene pair. The tools report predictions for only a subset of the genes in a given genome, which leaves many genes without an assigned label. We decided to label the genes with missing predictions as non-operonic, since leaving them out of the performance measures would drastically diminish the tools’ scores. This way, the metrics involving true negatives reported for the tools are the highest they could attain, assuming they would predict all the non-operonic gene pairs correctly. In a good faith attempt to replicate the work done by ProOpDB, we fed a neural network with the same architecture mentioned in their paper the same features (i.e, the inter-genic distance and STRING score for consecutive pairs of genes). We used our training dataset, which we believe to be richer, considering that they were training on only one organism (E. coli or B. subtilis). We fine tuned different hyper parameters, like the learning rate and the number of epochs, and experimented with different activation functions before settling on the one mentioned in their paper. Instead of generating STRING-like scores for the genes that miss that feature, we followed our previous approach of using a default value of 0.1. It is worth mentioning that the STRING database has been updated since the time of their first publication, and the current version has score assignments for most of the operonic gene pairs (> 99% of the pairs used in our training and testing datasets). Following this approach, we achieved a slightly lower true positive rate, with a slightly higher precision, leading to the same F1 score previously reported for their tool, and a slightly higher specificity, that is still the lowest across the three tools we used in our comparison.

We point out some of the challenges that undermine operon prediction in general. One limitation that faces predictors that rely on features such as gene conservation or functional assignment is the requirement to have such information about all the genes in a genome. So while such predictors might perform well on gene pairs that include the necessary features, their performance might drop considerably when making predictions over the entire genome^2^. Moreover, even though most methods validate their results by comparing their predictions over experimentally verified operons, the fact that the experimentally verified datasets are only available for a small subset of the sequenced genomes and that the datasets used vary between studies poses extra challenges making the comparison between the available tools non-straightforward. Brouwer et al tried to compare several methods using a uniform dataset and noticed a significant gap between the measures achieved and those reported in their original papers. The drop in performance was even higher when considering full operon predictions rather than separate gene pair predictions^29,32^. Finally, Some methods include the testing dataset as part of the training dataset, which leads to a reported accuracy that is significantly higher than what would be otherwise, given that the flow of information taking place in the training process would not be easily and readily transferable to novel genomes used as a testing dataset^4^. Some methods report a decrease in accuracy when the training and testing datasets belong to different organisms. In fact, the accuracy reported by many of the available tools drop anywhere between 11 and 30% when the training data and the operon predictions correspond to different organisms^4,32^. This lack of generalization places severe limitations on the methods’ applicability, especially when treating novel genomes. Even for known genomes, this poses the additional challenge of requiring more ground-truth data for operons in other genomes that belong to the same organism. For example, as mentioned earlier, Door switches between a linear model and a more complex decision-tree based model depending on the availability of the experimentally verified operons in the organism of the query genome that can be used for training the model. Our approach alleviates these challenges and generalizes successfully across organisms. As Table 4 shows, the genomes constituting our training dataset span different genera, but the performance of Operon Hunter when predicting operons in both E. coli and B. subtilis does not vary significantly. This is due to the fact that in the compare region viewer service that we use to construct the images, while the genomes aligned with the query genome are chosen based on evolutionary distance, they are eventually represented as images using a more generic method, that captures all underlying relevant operon features.

Much like feature engineering methods, casting tabular data to images encodes information in a way more amenable to learning without explicitly adding information. It can also be easily integrated with other data modalities in the latent vector representation to prevent information loss. We hypothesize this emerging trend of representing data with images will continue until model tuning and large-scale pre-training in scientific domains start to catch up with those in computer vision. Applications of this method are especially useful when the features are not easily quantifiable, as is the case in any application involving comparative genomics. Due to the generic nature of the visualizations and the available data augmentation techniques, we expect applications similar to our method to transform genomics problems where ground truth datasets are limited.

## 5 Author contributions statement

R.A. carried out the implementation and wrote the manuscript. R.S. and F.X. were involved in planning and supervised the work. All authors aided in interpreting the results. All authors provided critical feedback and commented on the manuscript.

## 7 Availability of data and materials

The datasets generated and/or analyzed during the current study are available in the GitHub repository, https://github.com/ridassaf/OperonHunter

## 8 Additional information

The authors declare that they have no competing interests.

